# Expansin couples salinity sensing to plant cell growth regulation

**DOI:** 10.64898/2026.11.14.766679

**Authors:** Ke Zhou, Daniel J. Cosgrove

## Abstract

In living plant cells, salt affects cell wall architecture directly and triggers diverse cellular responses that, in turn, feed back on wall organization and deposition. These intertwined effects obscure whether salt directly alters wall mechanical properties and thereby influences turgor-driven wall enlargement, the process underlying cell growth. In this study, we studied the tensile mechanical behavior of isolated cell wall materials at physiologically relevant salt concentrations. NaCl induced a small, reversible deformation without significantly altering the apparent axial tensile stiffness of hypocotyl walls. However, salt directly and reversibly inhibited acid-induced wall creep. Similar inhibition was observed in distinct cell wall types and in living Arabidopsis hypocotyls. Assays combining α-expansin extracts with cellulose filter paper further supported direct inhibition of wall-loosening activity by salt. Molecular dynamics simulations suggested a mechanism in which Na⁺ electrostatically perturbs the conserved active-site Asp and its associated hydrogen-bond network. Together, these findings indicate that salt directly inhibits acid growth by suppressing α-expansin-mediated wall creep, highlighting expansin as a physicochemical transducer linking environmental salinity to growth regulation.

## Introduction

Plant cell growth involves turgor-driven irreversible enlargement of primary cell walls, a process that overcomes mechanical resistance imposed mainly by the noncovalent network of cellulose microfibrils(*1*). To rapidly regulate this process, plants evolved a group of conserved, non-enzymatic proteins known as expansin(*2-4*). Expansins induce cell wall extension (creep) driven by turgor, potentially by disrupting the noncovalent binding between cellulose microfibrils, in a pH-dependent manner(*4-6*). This expansin-induced creep at low pH forms the basis of the canonical “acid growth” theory, whereby hormonal and environmental cues control growth partially by modulating this process(*7, 8*). Soil salinity is one of the most pervasive abiotic stresses in modern agriculture, severely restricting the growth and survival of terrestrial glycophytes, including most crop species(*9-11*). Early studies reported that divalent and trivalent cations(*12*), as well as high concentrations of NaCl(*13*), inhibit acid growth in isolated cucumber hypocotyl walls. However, whether salt acts on wall polymers, expansin proteins, or both - and how these interactions suppress wall extensions - remain unresolved. The physiological relevance of this inhibition also remains unclear.

Salt can rapidly and reversibly inhibit plant cell expansion, while prolonged salt exposure can cause irreversible growth suppression(*14*). Beyond immediate reduction in turgor, salinity is perceived through multiple pathways, including GIPC-dependent signaling at plasma membrane(*15*), the cytoplasmic SOS pathway(*16*), the secondary responses to oxidative stress(*14*), and perturbation of pectin structure(*17*). These responses can modify the apoplastic environment and transcriptionally govern cell-wall biosynthesis and deposition(*18*), thereby feeding back on wall mechanics and extension. Of particular relevance is the direct disruption of Ca^2+^-pectate crosslinking by salt, which softens the pectic matrix mechanically and triggers intracellular signaling(*17, 19, 20*). However, pectin softening, whether assessed in isolated pectin systems or by indentation of intact walls perpendicular to the growth axis, does not necessarily reflect changes in the tensile mechanical properties relevant to sustained wall extension along that axis(*21*). This distinction is consistent with recent work showing that noncovalent interactions between cellulose microfibrils provide the principal resistance to tensile stress(*22*) and that sliding between cellulose microfibrils underlies irreversible wall extension(*23*). It therefore remains unclear whether salt-induced structural changes alter wall mechanical properties in ways that directly influence wall extension and cell growth.

Here, we used complementary mechanical assays on isolated cucumber hypocotyl and maize coleoptile walls and on pure cellulose systems to examine how salt affects tensile mechanical properties independently of protoplast-mediated responses and changes in turgor(*4, 6*). We further combined molecular dynamics simulations of cucumber α-expansin CsEXPA1 with creep assays at different pH values to explore how cations might inhibit expansin activity through interactions with the conserved active-site Asp and its associated hydrogen-bond network.

## Results

### NaCl induces reversible hypocotyl deformation electrostatically

After freeze-thawing and flattening to remove cellular processes and turgor, apical segments from etiolated cucumber hypocotyls were clamped on both ends, soaked in an adjustable buffer, and subjected to a uniaxial constant load that replaces the axial component of cell turgor(*24*) (**Fig. 1A-1B**). This setup mimics hypocotyl elongation, restricting the key experimental variables to the composition of the controllable buffer. We previously utilized this system to characterize expansin’s cell wall creep activity(*4, 5, 12*). In neutral buffer (20 mM HEPES buffer, pH 7), the hypocotyl underwent minimal, time-dependent, and irreversible elongation (creep, red in **Fig. 1C-1D**), which reflects the gradual, permanent yielding of the cell wall polymers as they slide past one another(*22, 25*). Substituting the neutral buffer with an acidic buffer (50 mM MES, pH 5.2), a pH commonly found in the apoplast of growing primary cell walls, mimicked the typical apoplastic acidification that occurs in growing tissues(*26*), eliciting the characteristic acid-induced creep process: an abrupt acceleration of wall extension followed by deceleration to a new, stable, yet significantly higher rate(*4*) (blue in **Fig. 1C-1D**). In contrast, adding NaCl to the neutral buffer induced prompt but transient elongation without sustained effects (**Fig. 1C-1E**). This elongation is concentration-dependent (**Fig. 1F**), and fully reversible, as washing out the salt caused shrinkage that offset the initial elongation (**Fig. 1G, 1H, and Fig. S1**). Adding 100 mM sodium acetate (NaAc, adjusted to pH 7) induced an identical, reversible deformation, suggesting that the observed deformation may result from Na^+^ rather than Cl^-^ (**Fig. S2**). These results indicate that environmental Na^+^ does not cause the cell wall to undergo creep; instead, it results in an immediate and reversible wall deformation, distinct from the sustained and irreversible acid-induced extension.

**Fig. 1.**
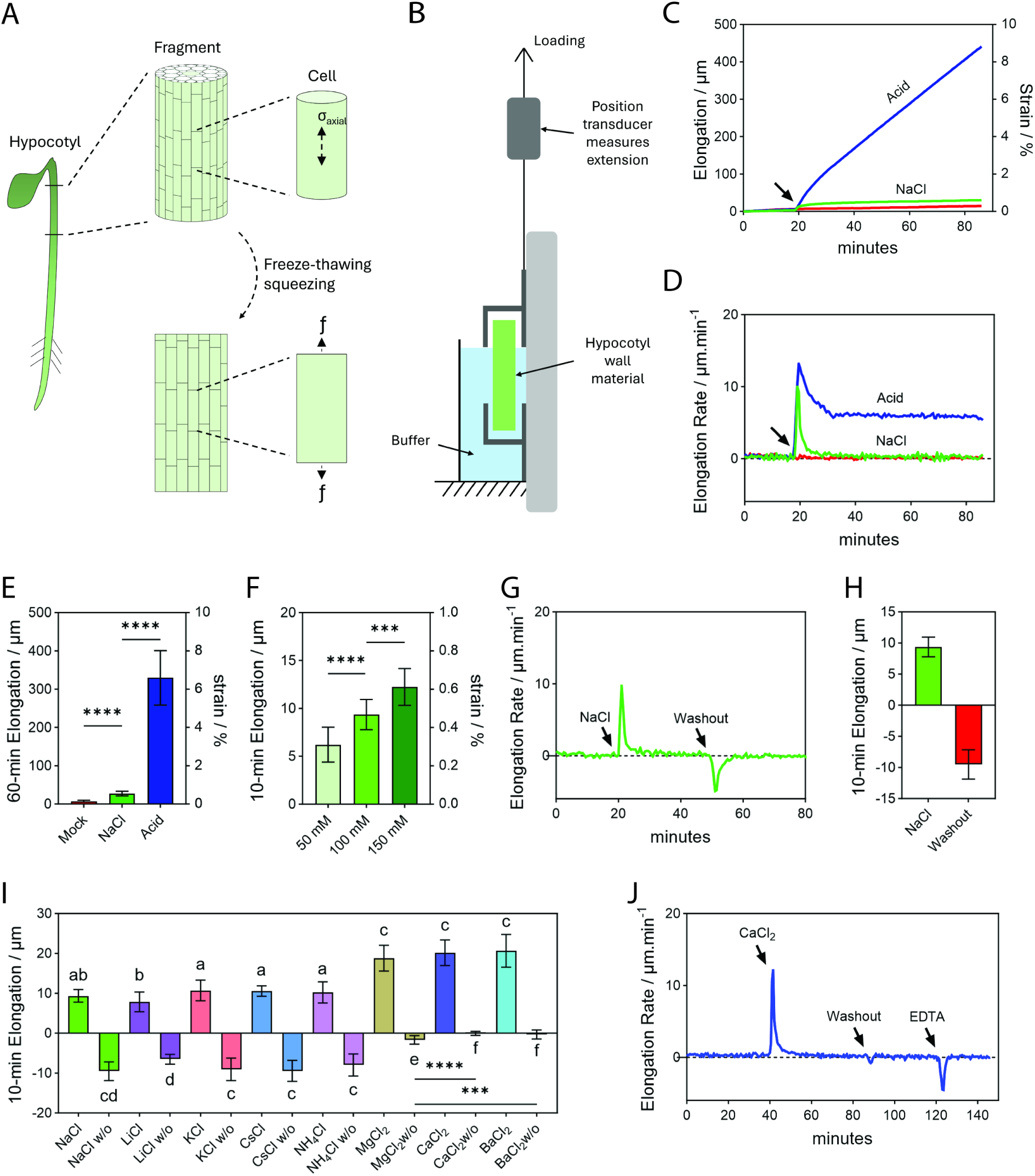
Acid- and NaCl-induced hypocotyl elongation *in vitro*. (**A**) Schematic of the *in vitro* hypocotyl elongation system. Actively elongating apical segments (0.5 cm) from three-day-old cucumber seedling hypocotyls were freeze-thawed and flattened to remove cell sap, then clamped at both ends and subjected to a 20 g uniaxial load. At cellular level, the axial component of turgor (σ-axial), which normally provides the driving force for elongation, is thus replaced by the applied load (f). (**B**) Recording of hypocotyl length every 30 s for determination of elongation and calculation of elongation rate. (**C**) Time course of hypocotyl elongation and (**D**) elongation rate in response to acid or NaCl. At 20 min (arrows), neutral buffer was replaced with either acidic buffer or neutral buffer containing 100 mM NaCl. Untreated, red; acid buffer, blue; 100 mM NaCl, green. (**E**) Total elongation during 60 min after treatment; means ± SD; n= 21, 21, 21. (**F**) Total elongation during 10 min in response to 50 mM, 100 mM, and 150 mM NaCl treatment in neutral buffer; means ± SD; n=14, 14, 12. (**G**) Representative elongation-rate trace for sequential addition and washout of 100 mM NaCl in neutral buffer. (**H**) Hypocotyl elongation after addition of 100 mM NaCl and shrinkage after washout; means ± SD; n=14. (**I**) Hypocotyl elongation during the 10 min after salt addition and shrinkage during the 10 min after washout (w/o) in response to 100 mM monovalent chloride salts and 50 mM divalent chloride salts; n = 12 (LiCl), 14 (NaCl), 12 (KCl), 13 (CsCl), 15 (NH₄Cl), 16 (MgCl₂), 16 (CaCl₂), and 16 (BaCl₂); means ± SD; different letters indicate statistically significant differences among treatments (P < 0.05, Student’s t-test). (**J**) Time course of elongation rate during sequential application of 50 mM CaCl_2_, washout, and 5 mM EDTA (arrows). The right y-axes of **C**, **E**, and **F** indicate the corresponding strain (%). Asterisks indicate significant differences (t-test; ***, p < 0.001; ****: p < 0.0001).

Next, we investigated the effects of other monovalent (LiCl, KCl, CsCl, and NH_4_Cl) and divalent (MgCl_2_, CaCl_2_, and BaCl_2_) chloride salts. All tested monovalent cations induced similar reversible deformations, except for LiCl, which caused a slightly weaker effect (**Fig. 1I**). Divalent cations elicited a more pronounced but largely irreversible deformation (**Fig. 1I**), which could be partially reversed by the chelating reagent EDTA (5 mM) (**Fig. 1J and Fig. S3**). This dependence on ionic valence, together with the partial reversal by EDTA, suggests that electrostatic interactions are largely responsible for the deformation.

Carboxyl groups, predominantly provided by the de-esterified polygalacturonic acid backbone of pectin, and supplemented by structural proteins(*27*), constitute the major electrostatic binding sites for metal ions in the primary cell wall. Among these electrostatic interactions, the Ca^2+^-bridged pectate “egg-box” structure is a particularly specialized form that is a well-known target of salt(*17, 19*). However, our data show that Mg^2+^, which has a minimal capacity to form this specific, stable configuration(*28*), caused an similar deformation that is nonetheless partly reversible (**Fig. 1I**). These observations therefore suggest that salt-induced hypocotyl deformation is dominated by generic electrostatic interactions with anionic sites in the cell wall, instead of arising from the specific Ca^2+^-pectate “egg-box” crosslinking interaction(*29*).

### NaCl insignificantly affects tensile mechanical properties of hypocotyl walls

Under constant load, salt induced reversible hypocotyl deformation but did not promote sustained, irreversible extension (creep). To complement these time-dependent measurements, we performed constant-rate tensile tests on freeze-thawed hypocotyls using the same initial gauge length (5 mm) as in the creep assays. Specimens were stretched at a constant crosshead speed of 3 mm/min while force and extension were recorded simultaneously. The resulting force-strain curves characterize the tensile mechanical response of the specimens by showing how force varies with strain. These curves were similar in the presence and absence of 100 mM NaCl in neutral HEPES buffer (**Fig. S4A)**. Their local slopes, which provide estimates of apparent tensile stiffness at a given strain, were also comparable over the common strain range (**Fig. S4B**). Together, these results suggest that NaCl may not substantially alter the apparent axial tensile stiffness of hypocotyl walls under the conditions tested.

### NaCl reversibly inhibits acid-induced hypocotyl elongation *in vitro*

We then applied NaCl (10-100mM), spanning mild to severe soil salinity in nature(*9*), to hypocotyl segments undergoing sustained acid-induced elongation. Adding salt caused a pronounced, concentration-dependent suppression of the elongation rate (**Fig. 2A-2B**). The inhibition was fully reversed upon washing out NaCl (**Fig. 2A**, **2C**), indicating that NaCl acts as a potent yet reversible inhibitor of acid-induced elongation. We tested additional mono- and divalent chloride salts, which produced responses that closely paralleled their effects on hypocotyl deformation in neutral buffer: all monovalent cations (35 mM) induced almost identical and reversible inhibition, with LiCl again an exception, showing a slightly attenuated effect (**Fig. 2D**); divalent salts (MgCl_2_, CaCl_2_, and BaCl_2_) at equimolar chloride concentrations (17.5 mM) caused a more potent inhibition that was largely irreversible; MgCl_2_ caused an inhibition comparable in potency to that of other divalent cations, yet this effect was partially reversible (**Fig. 2E**); and EDTA partially reversed the irreversible inhibition caused by Ca^2+^ (**Fig. 2F-2G**). Likewise, these results are consistent with a predominant contribution of electrostatic interactions, with little evidence for a significant role of Ca^2+^-pectate crosslinking for the inhibition.

**Fig. 2.**
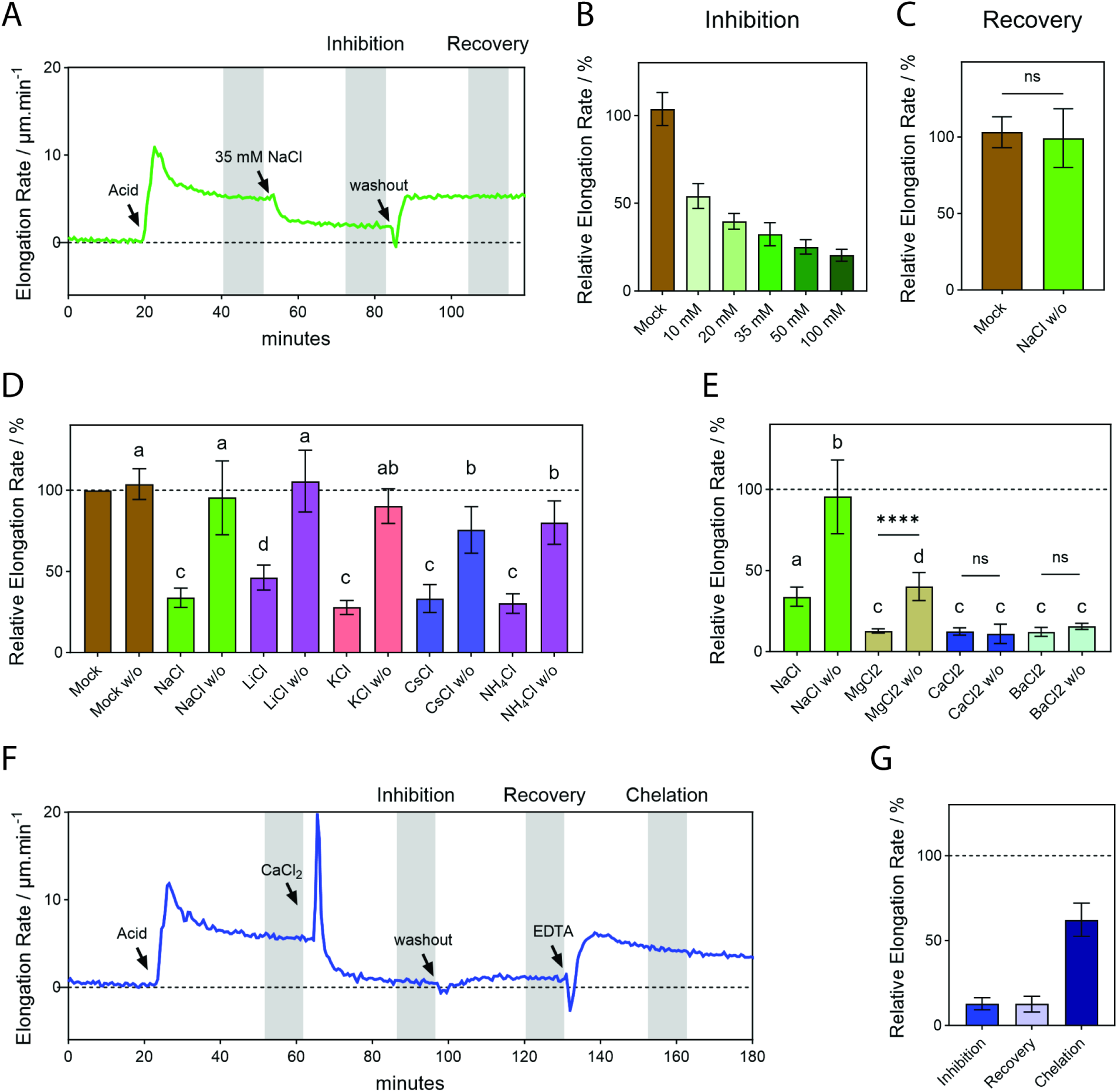
Inhibition of sustained acid-induced hypocotyl elongation by mono- and divalent chloride salts. (**A**) Representative elongation-rate trace for a hypocotyl segment during sustained acid-induced elongation in pH5.2 buffer, 35 mM NaCl treatment, and subsequent washout. Gray shading, time windows for quantification of inhibition and recovery in Figure B and C. (**B**) Relative elongation rate during the inhibition phase at the indicated NaCl concentrations (10-100 mM), normalized to the steady acid-induced rate before salt addition: means ± SD; n = 8 (mock), 21 (10 mM), 7 (20 mM), 14 (35 mM), 7 (50 mM), and 7 (100 mM). (**C**) Relative elongation rate during the recovery phase after washout of 35 mM NaCl compared with mock-treated controls; means ± SD; n = 7 (mock), 11 (NaCl); t-test; ns, not significant, P>0.05). (**D, E**) Relative elongation rate during salt treatment (35 mM monovalent chloride salts, or 17.5 mM divalent chloride salts) and after washout (w/o), normalized to the steady acid-induced rate before salt addition (dashed line, 100%). Data are means ± SD; n = 12 (mock), 12 (NaCl), 14 (LiCl), 11 (KCl), 7 (CsCl), 8 (NH₄Cl), 12 (MgCl₂), 7 (CaCl₂), and 7 (BaCl₂); means ± SD; t-test; ****: p < 0.0001; ns, not significant, P>0.05; different letters indicate statistically significant differences among treatments (P < 0.05, Student’s *t*-test). (**F**) Representative elongation-rate trace for a hypocotyl segment during sequential treatment with 17.5 mM CaCl₂, washout, and 5 mM EDTA; gray shading marks inhibition, recovery, and chelation periods used for quantification. (**G**) Relative elongation rates during the inhibition, recovery, and chelation phases in (F), normalized to the pre-salt acid-induced rate (dashed line, 100%); means ± SD; n=8.

### NaCl may directly suppress cellulose microfibril-loosening by *α*-expansin

Acid-induced elongation in cucumber hypocotyls is mediated by the pH-dependent action of expansins which may be extracted from the wall by NaCl(*30*). Notably, the first-isolated expansins, CsEXPA1 and CsEXPA2, were characterized from this NaCl-eluted protein extract(*4, 5*). In this study, we heat-inactivated wall specimens and then reconstituted acid-induced elongation by addition of NaCl-eluted wall proteins from cucumber hypocotyls (**Fig. 3A**). The reconstituted acid-induced elongation is amplified by the removal of Na^+^ and NH_4_^+^ present in the extract (see "Materials and Methods")(*4, 5*). NaCl (added at 90 min) potently inhibited this reconstituted elongation, and this inhibition was reversed upon salt washout (at 120 min) (**Fig. 3A-3C**). These results show that NaCl reversibly suppresses the reconstituted wall-loosening response, consistent with a direct, reversible inhibition of α-expansin-mediated loosening rather than irreversible damage or loss.

**Fig. 3.**
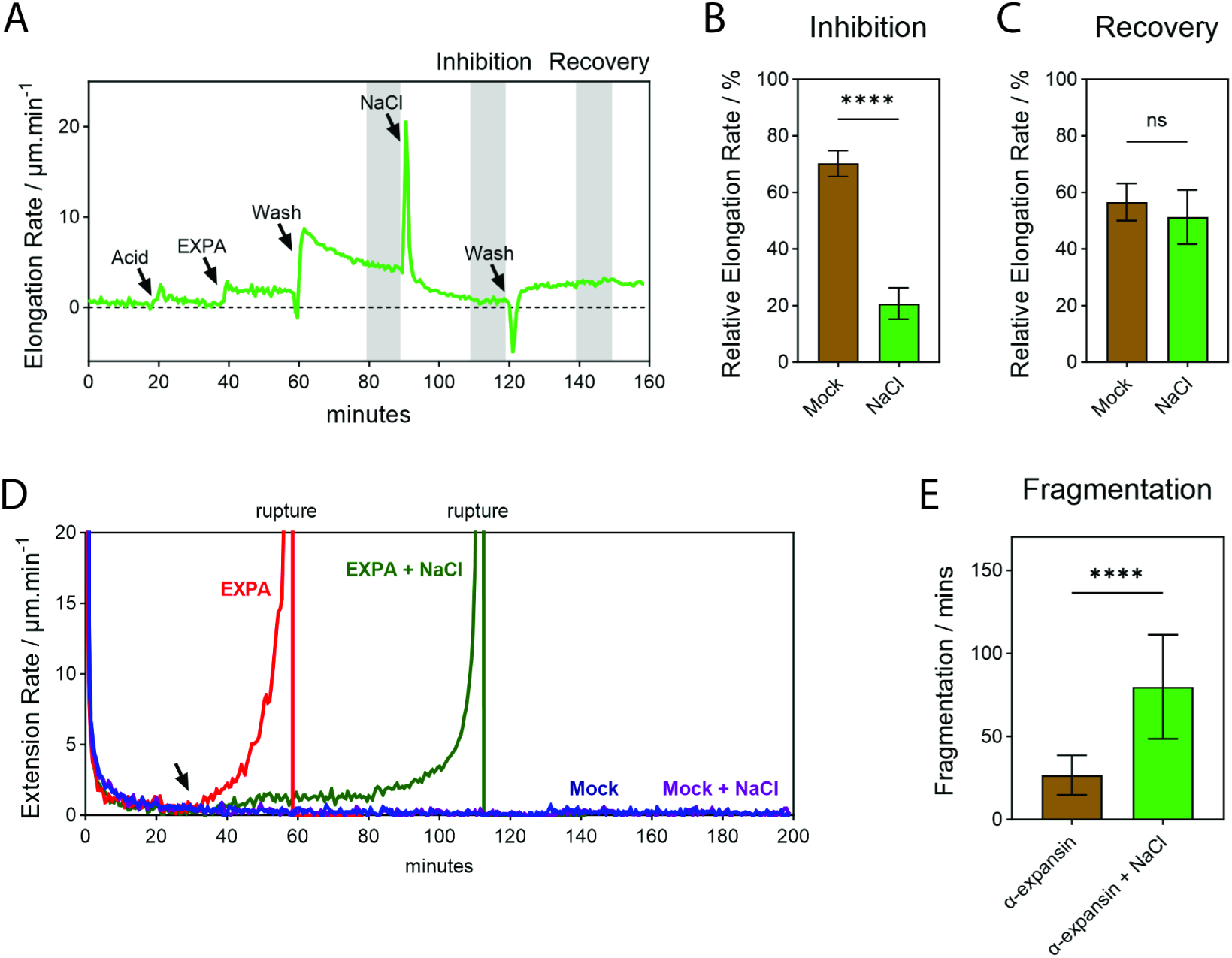
Salt-suppressed crude *α*-expansin-induced creep of hypocotyl segments and pure cellulose filter paper strips. (**A**) Representative elongation-rate trace of reconstituted acid-induced elongation and its salt-induced inhibition in a heat-inactivated hypocotyl segment. The neutral buffer was replaced with acidic buffer at 20 min, followed by addition of crude α-expansin extract in acidic buffer at 40 min. The extract was then washed out with acidic buffer at 60 min, 100 mM NaCl was added at 90 min, and washed out at 120 min (arrows). Gray shading indicates the inhibition and recovery windows used for quantification. (**B**, **C**) Relative elongation rate during the (B) inhibition and (C) recovery phases in mock and NaCl-treated samples, normalized to the crude α-expansin-induced rate before NaCl addition. Means ± SD; n = 10 (mock), 13 (NaCl); t-test; ****, P < 0.0001; ns, not significant, P>0.05. (**D**) Representative elongation-rate traces during creep of cellulosic filter paper strips. Treatments of crude α-expansin in the absence (red) or presence (green) of 100 mM NaCl, and acidic buffer in the absence (blue) or presence (purple) of 100 mM NaCl. Strips were pre-equilibrated in acidic buffer containing the corresponding NaCl concentration, and the arrow at 30 min indicates the time point of treatment. (**E**) Time from treatments to strip failure in the absence (n = 12) or presence (n = 10) of 100 mM NaCl; means ± SD; t-test; ****: p < 0.0001.

Given the difficulties of purifying EXPA proteins and the compositional complexity of native plant cell walls, we further minimized the system by replacing the cucumber hypocotyl with pure cellulose filter paper strips. Under constant tensile stress in an aqueous environment, the mechanical integrity of this material depends on noncovalent binding between cellulose fibers, analogous to those that stabilize the native cell wall(*31*). Previous work has established that exogenous α-expansin weakens these interactions, leading to unstable extension and mechanical failure of the filter paper(*5*). With a 12-gram load, hydrated filter paper strips (2.1 mm x 5 mm, Whatman 3# pure cellulose square filter paper) in acidic buffer (with and without 100 mM NaCl, but without added protein) exhibited negligible creep, without failing within the maximum observation period of 200 minutes) (**Fig. 3D, S5A-S5B**). The addition of NaCl-eluted cell wall proteins from cucumber hypocotyls, however, markedly accelerated filter paper extension, driving the strips into accelerating creep followed by failure. Additional 100 mM NaCl substantially suppressed the accelerated creep (**Fig. 3D, S5B**), prolonging the time-to-failure (**Fig. 3E, S6**). These findings in a pure-cellulose system therefore indicate that NaCl directly inhibits α-expansin-mediated disruption of load-bearing noncovalent interactions between cellulose microfibrils.

### The inhibition by salt is ubiquitous *in vitro* and *in planta*

Grass primary walls differ from dicot walls in matrix composition(*32*). In grasses the coleoptile has become the standard system for studying rapid growth and its hormonal regulation(*33-35*). It is a fast-elongating, hollow sheath surrounding the shoot apex and protecting the plumule; acid-induced elongation of coleoptile walls *in vitro* requires α-expansin activity(*6*). In the following experiments we substituted freeze-thawed maize coleoptiles for hypocotyls in our elongation assay and used this system to compare the effect of NaCl on freeze-thawed maize coleoptile walls undergoing acid-induced elongation. As shown in **Fig. 4A** and **4B**, salt suppressed acid-induced growth in coleoptile walls, suggested this NaCl effect may be a conserved feature of plant cell walls.

**Fig. 4.**
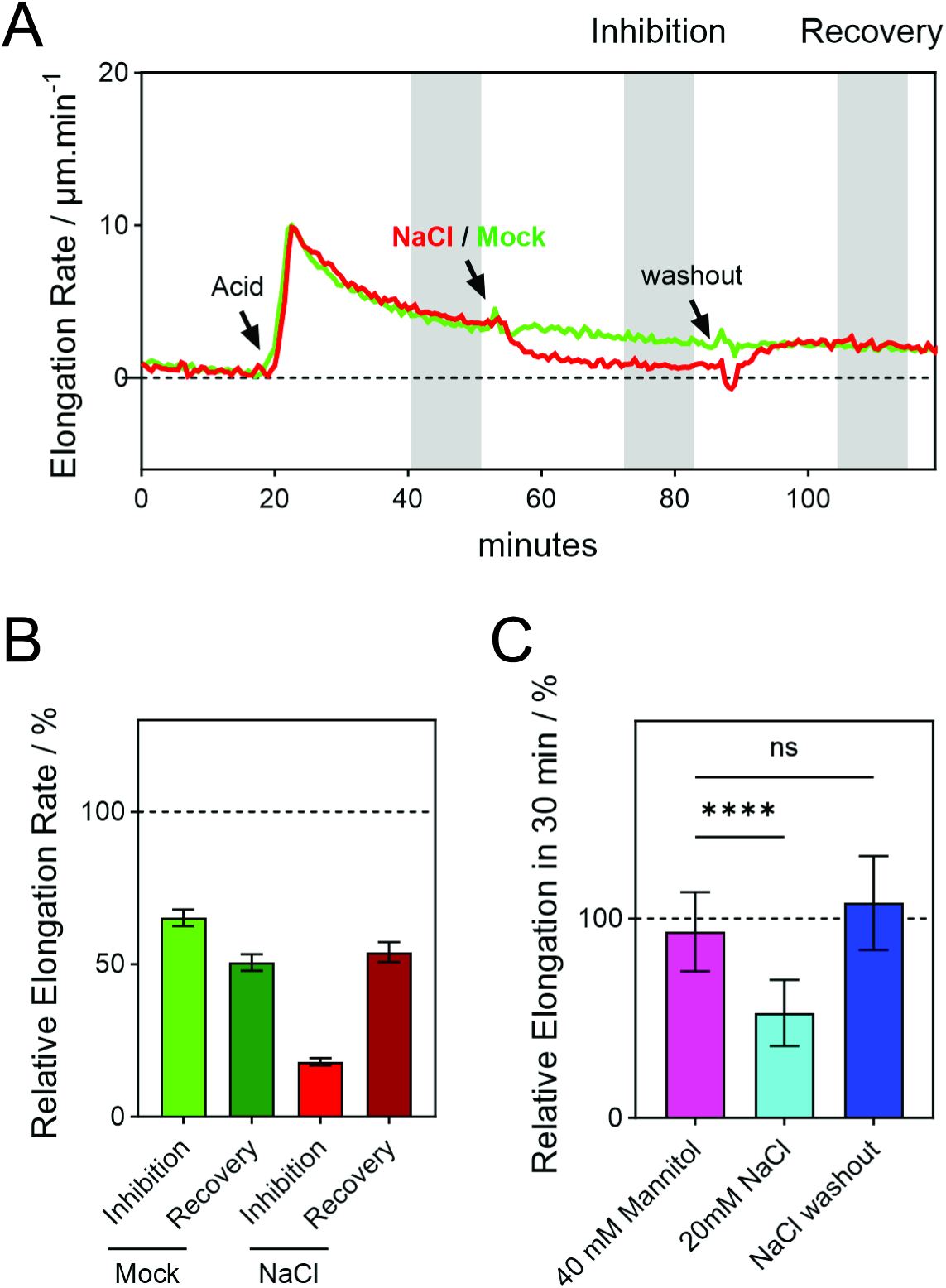
Salt-suppressed acid-induced elongation of coleoptile and live Arabidopsis hypocotyls. **(A**) Representative elongation-rate trace of freeze-thawed maize coleoptile during sustained acid-induced elongation in pH5.2 buffer, 35 mM NaCl treatment, and subsequent washout. Gray shading, time windows for quantification of inhibition and recovery in Figure B. (**B**) Relative elongation rate during the inhibition and recovery phases of maize coleoptiles, normalized to the steady acid-induced rate before salt addition: means ± SD; n = 8 (mock), 8 (NaCl). (**C**) Relative 30-min elongation of living Arabidopsis hypocotyls. Three-day-old seedlings were treated with 20 mM NaCl or 40mM mannitol in 50mM MES buffer (pH 5.2). The same 11 hypocotyls were measured during NaCl treatment and after washout; n = 10 for mannitol. Elongation was expressed as a percentage of the 30-min length increase in the untreated control (100%; dashed line). Data are means ± SD; t-tests for the comparisons indicated by brackets; ****, P < 0.0001; ns, not significant.

In living plant tissues, cell growth is governed by turgor-driven wall extension. Beyond reducing turgor, salt acts on walls via multiple ways, including direct disruption of Ca²⁺-pectate crosslinks(*17, 19, 20*), transcriptional remodeling wall biosynthesis and deposition(*18*), and alteration of apoplastic environment(*36*). To decouple the direct impact of salt on cell wall extension from these multifaceted and interacting processes, we incubated live Arabidopsis hypocotyls in 50 mM MES buffer (pH 5.2) to stabilize apoplastic pH. The observation period was limited to 30 minutes to minimize transcriptional effects. Application of 20 mM NaCl markedly but reversibly suppressed the elongation (**Fig. 4C**), whereas 40 mM mannitol, which imposes a comparable osmotic stress, had negligible effect (**Fig. 4C**). These *in vivo* results indicate that salt triggers rapid and reversible growth arrest beyond turgor reduction, consistent with salt-induced inhibition on α-expansin-mediated wall creep.

### A conserved H102-T11-D104 hydrogen-bond network in CsEXPA1

To investigate how salt interferes with α-expansin-induced wall creep, we performed molecular dynamics (MD) simulations using CsEXPA1, an ortholog of Arabidopsis EXPA8 and a major active expansin in NaCl-eluted cell-wall protein extracts from cucumber hypocotyls*(4, 5)*. Modeling by AlphaFold (AF-Q39625-F1-v6(*37, 38*)) shows a canonical expansin structure consisting of an N-terminal domain-1 with structural features resembling Glycosyl Hydrolase Family-45 (GH45) and a C-terminal domain-2 with features of carbohydrate-binding module-63 (CBM63)(*39-43*)(**Fig. 5A**). In GH45 enzymes, the conserved Asp of HFD motif acts as the catalytic proton donor, protonating the glycosidic oxygen by its side chain carboxyl group to promote β-1,4-glycosidic bond cleavage, and may subsequently be re-protonated through a conserved hydrogen-bond network involving a His-Thr-Asp proton relay(*44*)(**Fig. 5B**). The corresponding Asp is critical for wall-creep activity in both bacterial expansins(*41*) and plant EXPAs(*45*). Notably, although expansins lack hydrolytic activity and the complete proton-acceptor/proton-relay machinery of GH45 enzymes(*2*), canonical EXPAs retain the conserved Thr and HFD motif corresponding to T11 and H102-F103-D104 in CsEXPA1(*45, 46*)(**Fig. 5C** and **Fig. S7**).

**Fig. 5.**
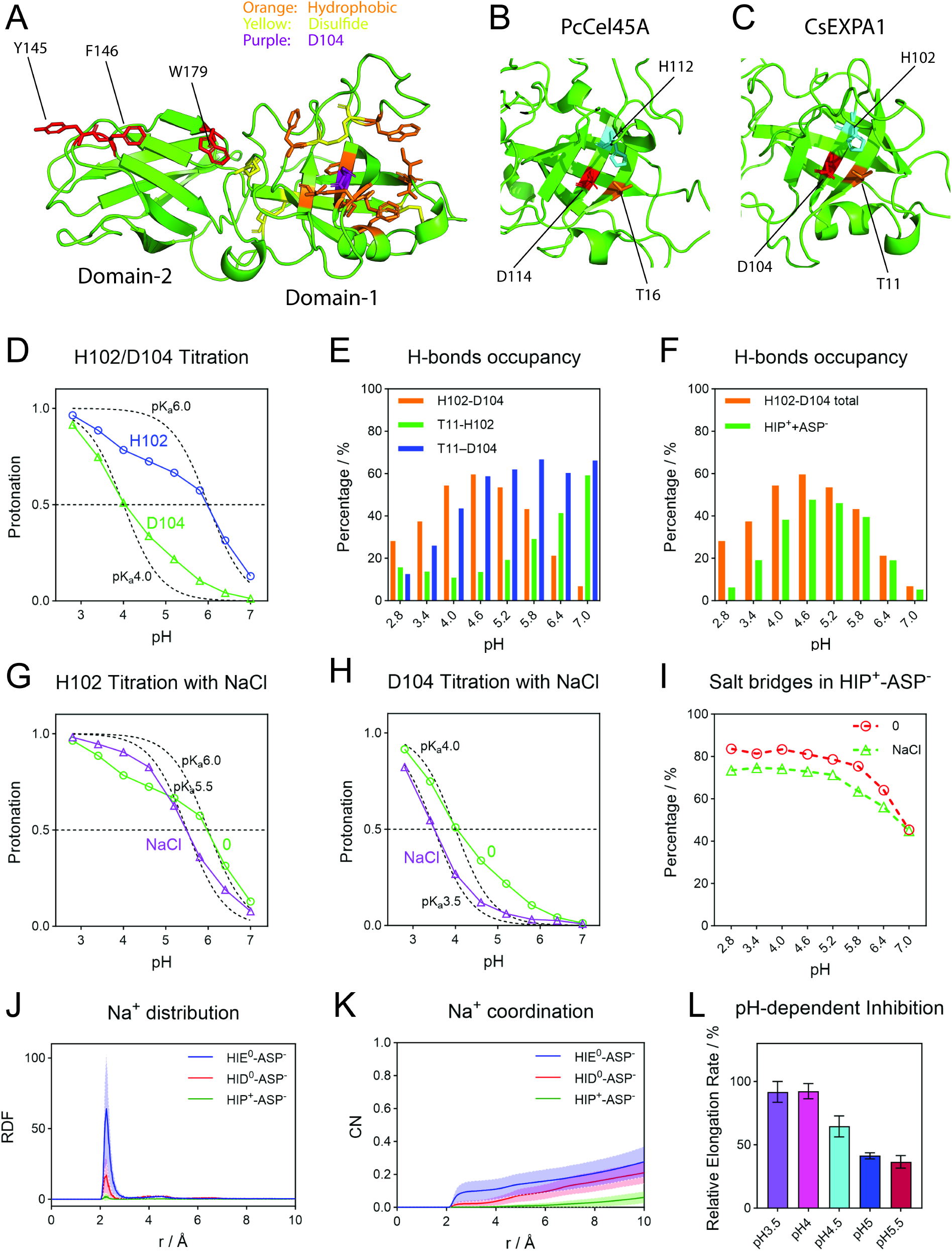
Molecular basis of pH- and salt-dependent regulation of CsEXPA1. (**A**) Predicted CsEXPA1 structure from Alphafold. Conserved aromatic residues in domain-2 (red), hydrophobic/aromatic residues lining the domain-1 groove (orange), D104 (purple) and predicted disulfide bonds (yellow) are shown. (**B, C**) Corresponding H-T-D residues in the GH45 enzyme PcCel45A (B; H112, T16 and D114) and CsEXPA1 (C; H102, T11 and D104). His, Thr and Asp are shown in cyan, orange and red, respectively. (**D**) pH-REMD titration curves of H102 (blue) and D104 (green) without added NaCl. (**E**) Hydrogen-bond occupancies for H102-D104 (orange), T11-H102 (green) and T11-D104 (blue). (**F**) Total H102-D104 hydrogen-bond occupancy (orange) and the contribution from the HIP⁺-ASP^-^ state (green). (**G, H**) Titration curves of H102 (G) and D104 (H) without added NaCl (green) or with 100 mM NaCl (purple). In D, G and H, symbols indicate simulated protonated fractions; black dashed curves show Henderson-Hasselbalch reference profiles at the indicated p*K*ₐ values. Horizontal dashed lines indicate a protonated fraction of 0.5. (**I**) H102-D104 salt-bridge occupancy within the HIP⁺-ASP^-^ state without added NaCl (red) or with 100 mM NaCl (green). Analyses in D-I were derived from pH-REMD simulations. (**J, K**) Na⁺ radial distribution functions (RDF; J) and cumulative coordination numbers (CN; K) around the D104 carboxylate oxygens in conventional MD simulations containing 100 mM NaCl. D104 was fixed as ASP⁻ and H102 as HIE⁰ (blue), HID⁰ (red) or HIP⁺ (green). r denotes the Na⁺-carboxylate O distance. Lines indicate means and shaded bands indicate ± SD across three independent 500-ns simulations. (**L**) pH-dependent inhibition of acid-induced hypocotyl elongation by Na⁺. Freeze-thawed hypocotyl segments under a 20-g load were transferred from neutral HEPES buffer to 20 mM sodium acetate buffer at the indicated pH, followed by the same buffer supplemented with 80 mM NaCl. Relative inhibition was calculated as in Fig.2A and 2B. Data are means ± SD; n = 8 per pH.

Given the proton-donating role of the homologous Asp in GH45 enzymes, we first sought to determine the protonation state of the D104 side-chain carboxyl group. We therefore characterized the pH-dependent protonation behavior of titratable residues in CsEXPA1 by constant-pH replica-exchange molecular dynamics (pH-REMD)(*47*). Conformations were sampled by MD in explicit solvent, and discrete protonation-state changes were attempted at fixed intervals using Monte Carlo moves evaluated with a generalized Born (GB) implicit-solvent model(*47*). Simulations spanned pH2.8-7.0 in 0.6-unit increments. The modified protocol reproduced experimentally determined p*K*ₐ values for titratable residues in hen egg-white lysozyme (HEWL)(*47, 48*)(**Table. S1**). We next analyzed the titration behavior of Asp, Glu and His residues in domain-1 of CsEXPA1 (**Table. S2**). D104 and the neighboring H102 exhibited unusually smooth titration curves that deviated markedly from Henderson-Hasselbalch behavior (**Fig. 5D**). T11, H102, and D104 formed a dynamic hydrogen-bond network through their hydroxyl, imidazole, and carboxyl groups, respectively (**Fig. 5E**). The T11A substitution, which removes hydroxyl group from the Thr, shifted the protonation equilibria of both H102 and D104 towards their neutral states (**Fig. S8A-S8B**), consistent with a role for T11 in stabilizing their charged states within the hydrophobic groove (**Fig. 5A**). Most H102-D104 hydrogen bonds were charge-assisted, forming between positively charged H102 (HIP⁺) and negatively charged D104 (ASP^-^) (**Fig. 5F**). Consistent with these strong interactions, fixing the protonation state of either residue altered the titration behavior of the other, indicating strong protonation coupling (**Fig. S8C-S8D**). Together, these simulations suggest that, as in GH45 enzymes, the active-site Asp D104 in domain-1 of CsEXPA1 lies within an H102-T11-D104 hydrogen-bond network. This network may modulate the protonation state of the D104 side chain, which could be important for CsEXPA1 activity.

### Potential mechanisms for salt-induced inhibition

Including 100 mM NaCl in the explicit solvent of our pH-REMD simulations markedly altered the titration behavior of H102 and D104, bringing both curves into closer agreement with the standard Henderson-Hasselbalch relation (**Fig. 5G-5H** and **Table. S3**), whereas the protonation behavior of most other Asp, Glu and His residues remained largely unchanged (**Fig. S9** and **Table. S3**). Most H102-D104 hydrogen bonds were charge-assisted contacts between protonated H102 (HIP⁺) and deprotonated D104 (ASP^-^) (**Fig. 5F**). NaCl reduced salt-bridge occupancy within this HIP⁺-ASP^-^ state (**Fig. 5I**), consistent with perturbation of electrostatic coupling between the two residues by the added ions(*49*). In parallel, overall T11-D104 hydrogen-bond occupancy (**Fig. S10**). Together, these changes suggest that NaCl may reorganize the H102-T11-D104 hydrogen-bond network, in part through electrostatic screening of H102-D104 interactions, potentially influencing CsEXPA1 activity.

In our pH-REMD simulations, protonation-state changes were evaluated using a GB model that excluded explicit ions(*47*) and therefore may not fully capture the coupling between Na⁺ coordination, D104 protonation and H102-T11-D104 hydrogen-bond network. We therefore performed conventional MD simulations in explicit solvent containing 100 mM NaCl, with D104 fixed as ASP^-^ and H102 as HIP⁺, HID⁰ or HIE⁰. Na⁺ transiently coordinated D104 when H102 was neutral, with a preference for the HIE⁰ state, whereas protonated H102 (HIP⁺) strongly suppressed this coordination (**Fig. 5J-5K** and **Fig. S11A**). A role for cation coordination in CsEXPA1 inhibition was further supported by our findings with divalent cations. In simulations of the HIE⁰-ASP^-^ state in explicit solvent containing 50 mM CaCl₂, Ca²⁺ coordination to the D104 carboxylate persisted after initial contact for the remainder of the trajectories (**Fig. S11B**). This sustained coordination is consistent with the stronger, less reversible inhibition of hypocotyl elongation by divalent cations (**Fig. 2E**) and the partial reversal of CaCl₂ inhibition by EDTA chelation (**Fig. 2D 2G**). Together, these findings suggest that cation coordination to the D104 side-chain carboxylate, which could alter the local electrostatic environment or perturb D104’s interactions with cellulose or neighboring residues, may contribute to salt inhibition of CsEXPA1.

Na⁺ coordination to the D104 side-chain carboxylate was strongly suppressed when H102 was positively charged (HIP⁺) (**Fig. 5J-5K**). We therefore predicted that the higher fraction of H102 in the HIP⁺ state at lower pH (**Fig. 5D**) would electrostatically disfavor Na⁺ coordination and thereby reduce Na⁺-mediated inhibition of CsEXPA1. Consistent with this prediction, inhibition of acid-induced hypocotyl elongation by 100 mM NaCl weakened progressively as pH decreased (**Fig. 5M**).

Together, these results support a model in which Na^+^ reversibly perturbs the H102-T11-D104 hydrogen-bond network and transiently coordinates with the active-site Asp D104, thereby inhibits its wall-loosening activity.

## Discussion

Cell wall-localized expansins can be activated by apoplastic acidification, facilitating wall creep and turgor-driven cell expansion(*4*). This pH-dependent process underpins acid-growth theory and provides a conceptual framework for understanding how hormonal and environmental signals regulate plant cell growth(*3, 25, 32*). This mechanism contributes to both diffuse growth*(4)* and tip growth(*45*). Our findings suggest that expansins, the central effectors of ‘acid growth’, can also function as physicochemical sensors of salinity. Salt-induced inhibition of their wall-loosening activity directly translates salt exposure into rapid, reversible growth inhibition. This direct physicochemical mechanism could operate independently of canonical signaling pathways, such as FERONIA-mediated cell wall integrity signaling and GIPC-dependent salt sensing that elicit Ca^2+^ signals(*15, 17*). Moreover, it also offers a mechanistic explanation for long-standing observations of the immediate and reversible inhibition of plant cell growth by salinity(*50, 51*).

Ca²⁺-pectate crosslinking is a specific electrostatic interaction: Ca^2+^ ions bridge de-esterified polygalacturonic acid chains to form the characteristic “egg-box” configuration(*52, 53*). High salinity can disrupt these Ca²⁺-mediated bridges, thereby softening the pectic matrix(*17, 54*). In our study, however, NaCl neither induced sustained wall creep nor significantly altered the apparent axial tensile stiffness of hypocotyl walls. Instead, our results suggest that NaCl suppresses wall expansion through direct, rapid, and reversible inhibition of expansin activity. These observations are consistent with our previous findings that softening of pectic wall components and growth-associated loosening are not necessarily coupled processes (*21, 55*). We therefore propose that salinity-induced changes in pectate crosslinking may influence cell growth primarily through cell wall integrity signaling and downstream responses(*17*), rather than through direct mechanical effects on wall expansion.

Although salinity imposes fundamental physicochemical constraints on plant cell growth, halophytes have adapted to chronically saline environments(*56*), and monocotyledonous seagrasses even thrive in full-strength seawater(*57*). Our findings raise the question of how plants circumvent this physicochemical constraint. Possible strategies include increasing EXPA abundance, consistent with reports that EXPA overexpression enhances salt tolerance(*18*); further acidifying the apoplast to alleviate inhibition; or deploying salt-adapted EXPA variants or other expansin family members. The unexpectedly broad functional diversity recently revealed within the expansin superfamily(*45*) suggests that such specialization is plausible. Defining these adaptive mechanisms could inform the rational breeding of crops with enhanced salt tolerance.

## Acknowledgments

We thank Jingyi Yu and Chinghai Li for initial discussions on early versions of this work, and C. Denise Okafor for helpful technical feedback on the molecular dynamics methods and analysis.

## Author contributions

Conceptualization: KZ

Methodology: KZ

Investigation: KZ

Visualization: KZ

Funding acquisition: DJC

Project administration: KZ

Supervision: KZ, DJC

Writing – original draft: KZ

Writing – review & editing: KZ, DJC

## Data, code, and materials availability

All data are available in the main text or the supplementary materials.

---

**Figure. S1.**
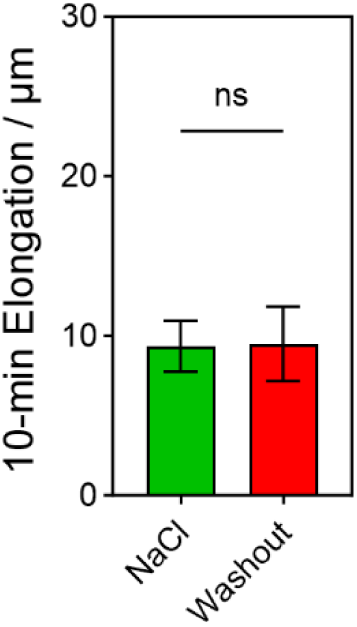
Absolute values of 10-minute hypocotyl elongation and shrinkage after adding and washing out 100 mM NaCl in the neutral buffer; means ± SD; n = 14; t-test; ns (not significant), p>0.05.

**Figure. S2.**
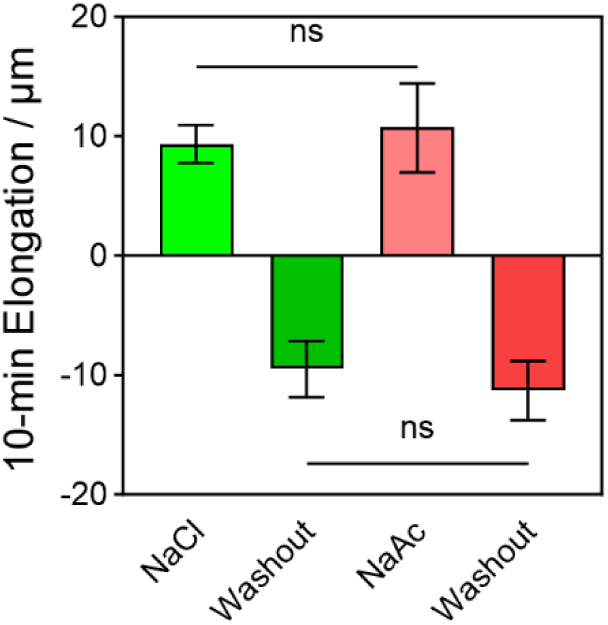
10-minute hypocotyl elongation and shrinkage during addition and washout of 100 mM NaCl or NaAc in neutral buffer. Data are means ± SD; n = 14 (NaCl) and 15 (NaAc); t-test; ns, P > 0.05.

**Figure. S3.**
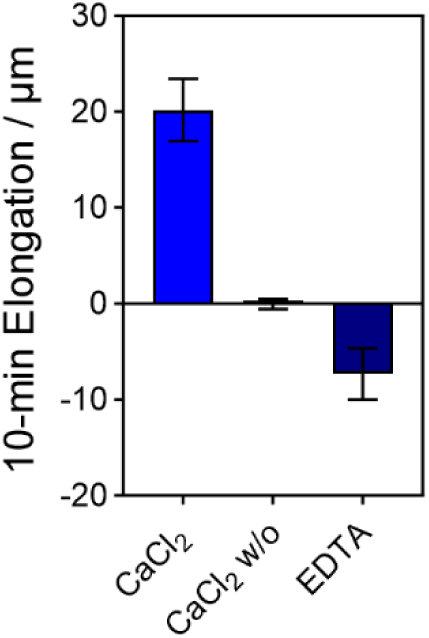
10-minute hypocotyl elongation and shrinkage during addition of 50 mM CaCl_2_, washout, and subsequent chelation with 5 mM EDTA in neutral buffer. Data are means ± SD; n = 16; t-test; ns (not significant), p>0.05.

**Figure S4.**
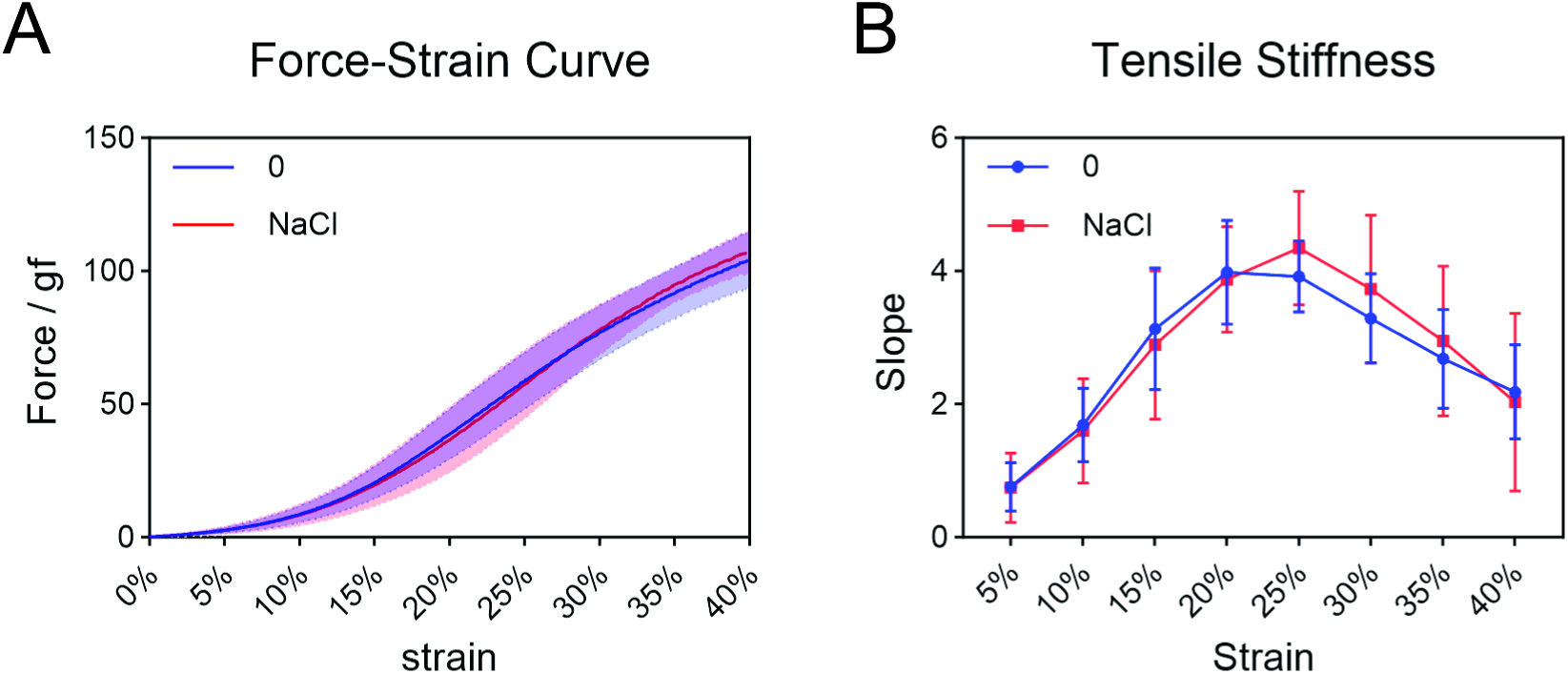
Effects of NaCl on the tensile mechanical properties of hypocotyl walls. (A) Force-strain curves of freeze-thawed cucumber hypocotyl wall specimens stretched along the growth axis in neutral HEPES buffer containing 0 (blue) or 100 mM NaCl (red). Specimens were stretched at a constant crosshead speed of 3 mm/min with an initial gauge length of 5 mm. Extension was referenced to the point at which the load first reached 0.1 gf. Lines show means, and shaded bands indicate ± SD. (**B**) Apparent tensile stiffness estimated from local slopes of the force-strain curves at 5%, 10%, 15%, 20%, 25%, 30%, 35%, and 40% strain. For each specimen, local slopes were estimated by linear regression within approximately ± 1 percentage point of each target strain. Slopes were expressed in gf per percentage point of strain. Symbols and error bars represent means ± SD. n = 19 control and 17 NaCl-treated hypocotyls.

**Figure S5.**
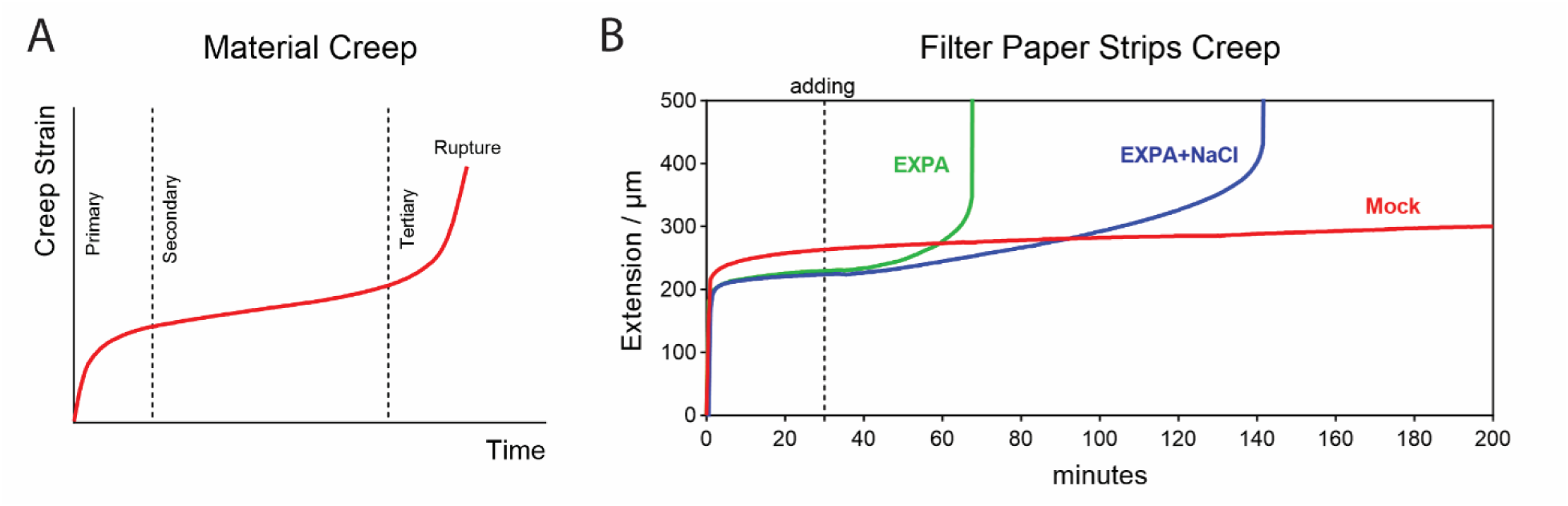
Creep of filter paper strips. (**A**) Schematic diagram of a typical material creep curve, which consists of three stages: primary (initial, characterized by a decreasing creep rate), secondary (steady-state, with an approximately constant creep rate), and tertiary (accelerated, leading to failure). (**B**) Representative extension curve of filter paper strips. Dry filter paper strips were clamped at both ends, hydrated with acidic buffer, and their length was recorded immediately thereafter. Crude EXPA extract solubilized in the pH 5.2 MES buffer without (green) and with (blue) 100 mM NaCl was applied at 30 min.

**Figure S6.**
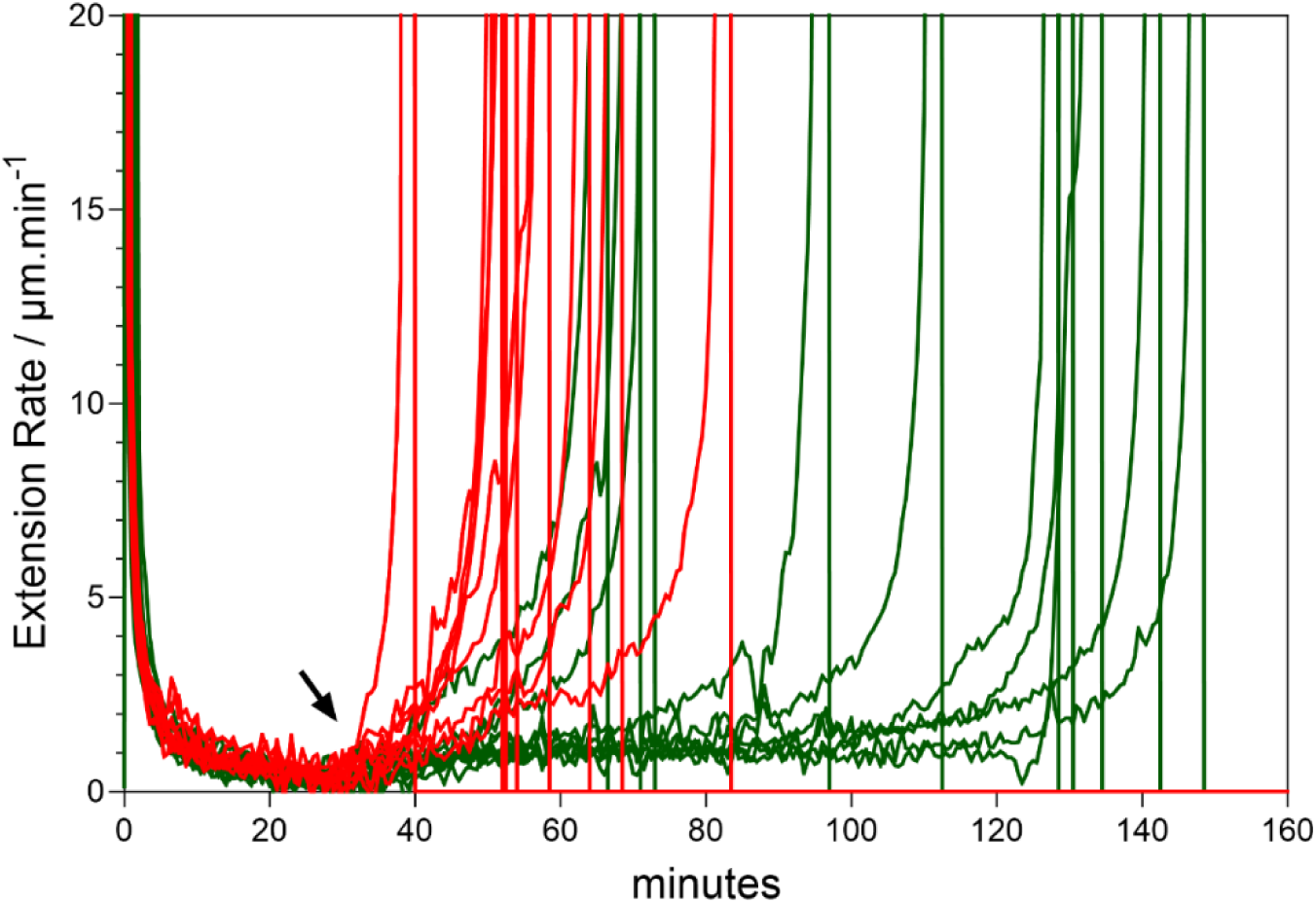
Extension rate of cellulose filter paper strips upon application of crude α-expansin with or without NaCl. Crude EXPA solubilized in acidic buffer without (red) or with (green) 100 mM NaCl was applied at 30 min (arrow) to filter paper strips pre-hydrated in acidic buffer containing the corresponding NaCl concentration. The y-axis shows extension rate; the sharp increase and subsequent drop of the rate to zero indicate failure of the strip. The average time to failure after expansin addition is summarized in Fig. 4e.

**Figure S7.**
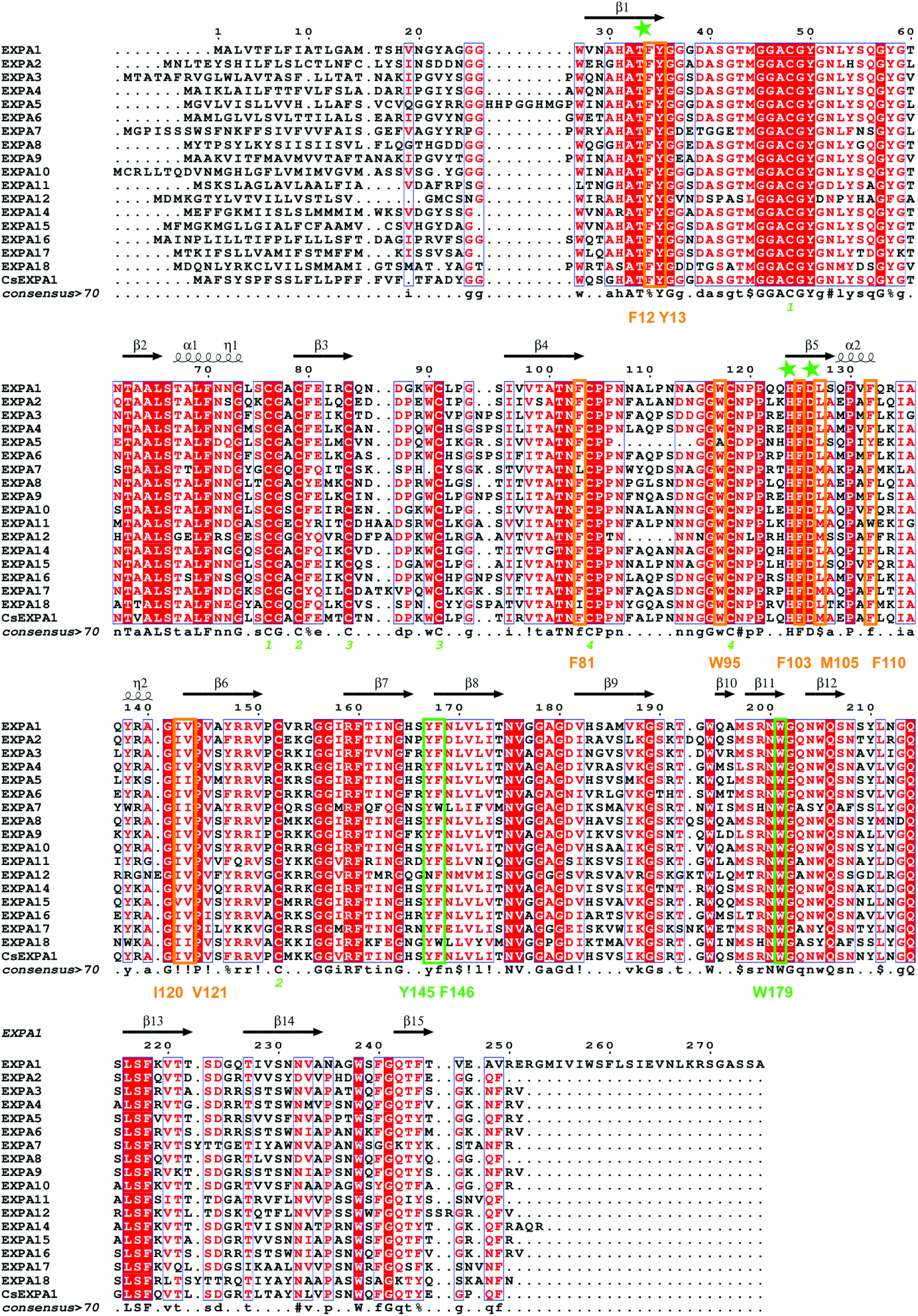
Sequence alignment of canonical Arabidopsis EXPA proteins and CsEXPA1. Conserved Thr, His and Asp residues (T-H-D) are marked with green stars. Conserved aromatic residues in domain-2 and conserved hydrophobic/aromatic residues in domain-1 are boxed in green and orange, respectively. The corresponding CsEXPA1 residues are labelled below the alignment in matching colors. Matching green numbers beneath conserved cysteine residues indicate predicted disulfide bonds.

**Figure S8.**
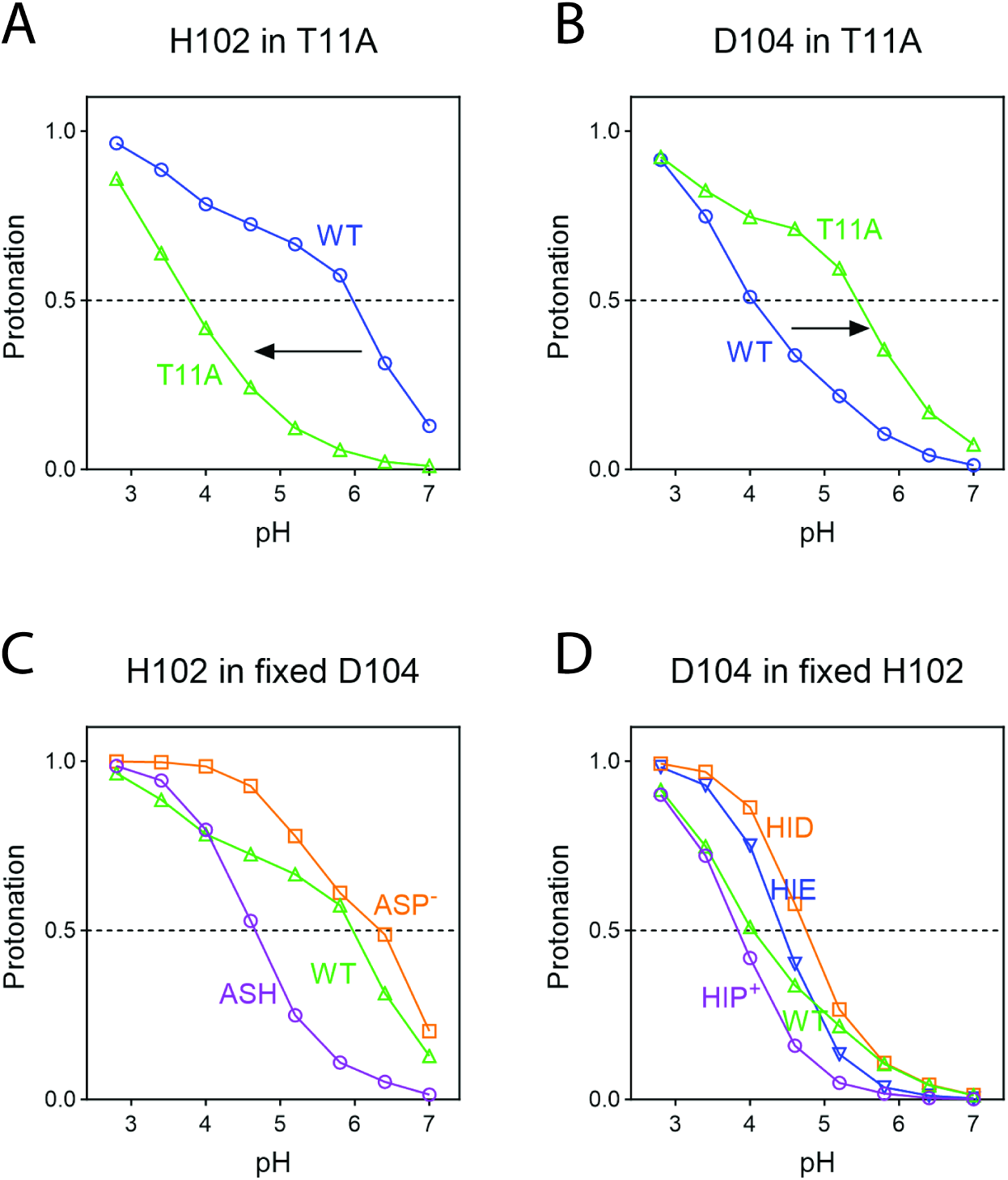
Protonation coupling within the H102-T11-D104 network of CsEXPA1. (**A, B**) pH-REMD titration curves of H102 (A) and D104 (B) in wild-type (WT; blue) and T11A (green) CsEXPA1. Arrows indicate shifts in the protonation transitions caused by T11A. (**C**) H102 titration with D104 fixed in its protonated, neutral state (ASH; purple) or deprotonated, negatively charged state (ASP^-^; orange). (**D**) D104 titration with H102 fixed as neutral HID^0^ (orange) or HIE^0^ (blue) tautomers, or in the doubly protonated, positively charged HIP⁺ state (purple). HID^0^ and HIE^0^ carry a proton on the N_δ_1 and N_ε_2 nitrogen atoms of the imidazole ring, respectively. Fixed protonation states are labelled in colors matching the corresponding curves. WT curves (green) are included for comparison in C and D. Symbols indicate simulated protonated fractions; dashed horizontal lines indicate a protonated fraction of 0.5.

**Figure S9.**
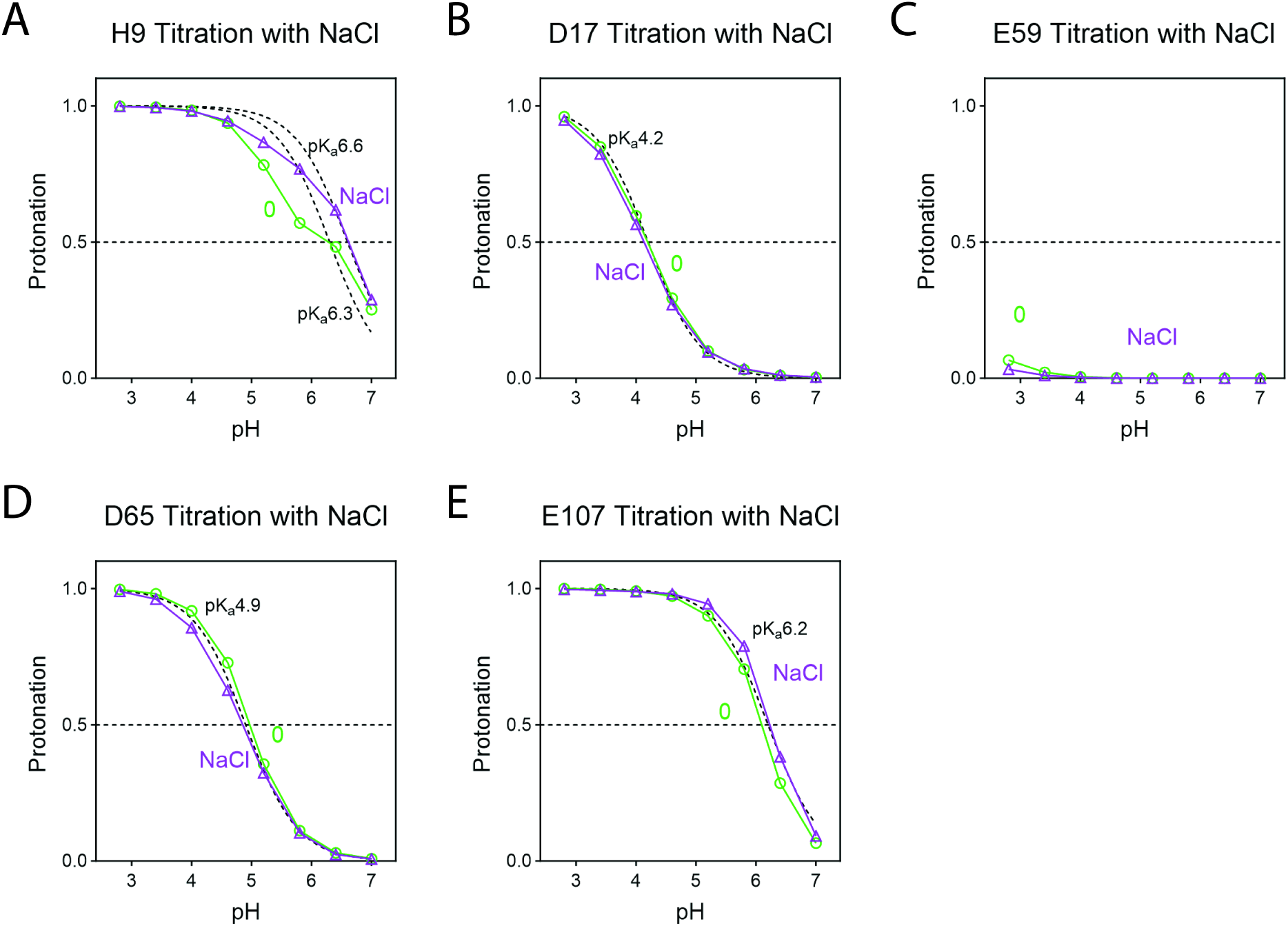
Effects of NaCl on the titration behavior of additional residues in domain-1 of CsEXPA1. Titration curves of (**A**) H9, (**B**) D17, (**C**) E59, (**D**) D65 and (**E**) E107, without added NaCl (green circles) or with 100 mM NaCl (purple triangles) in the explicit solvent. Points indicate protonated fractions obtained from pH-REMD simulations across pH 2.8-7.0. Black dashed curves show Henderson-Hasselbalch reference profiles at the indicated p*K*ₐ values. Horizontal dashed lines indicate a protonated fraction of 0.5.

**Figure S10.**
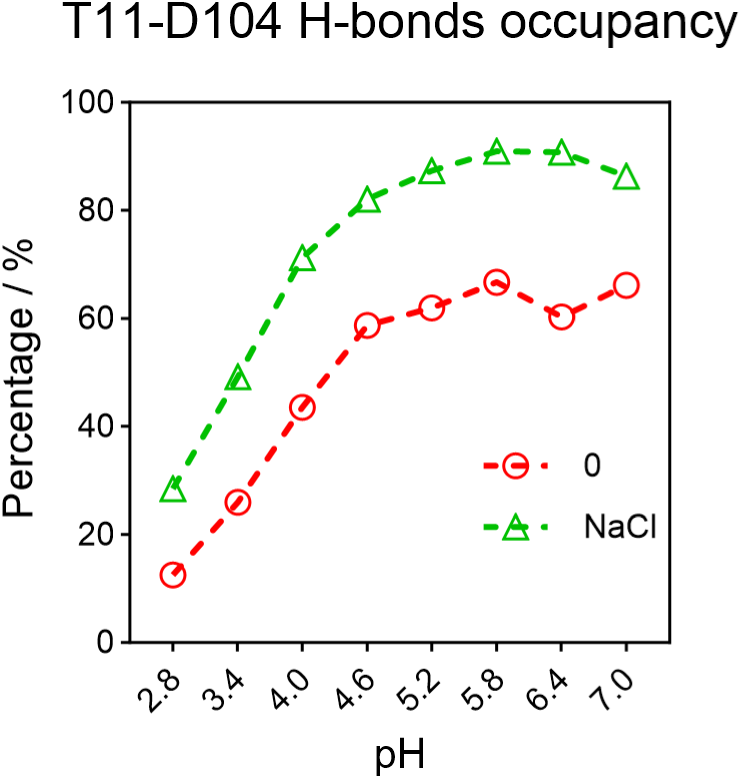
Effects of NaCl on T11-D104 hydrogen-bond occupancy in CsEXPA1. Hydrogen-bond occupancy between T11 and D104 was determined from pH-REMD simulations spanning pH 2.8-7.0, without added NaCl (red open circles) or with 100 mM NaCl (green open triangles) in the explicit solvent. Occupancy was calculated across all protonation states and expressed as a percentage of the total sampled frames at each pH.

**Figure S11.**
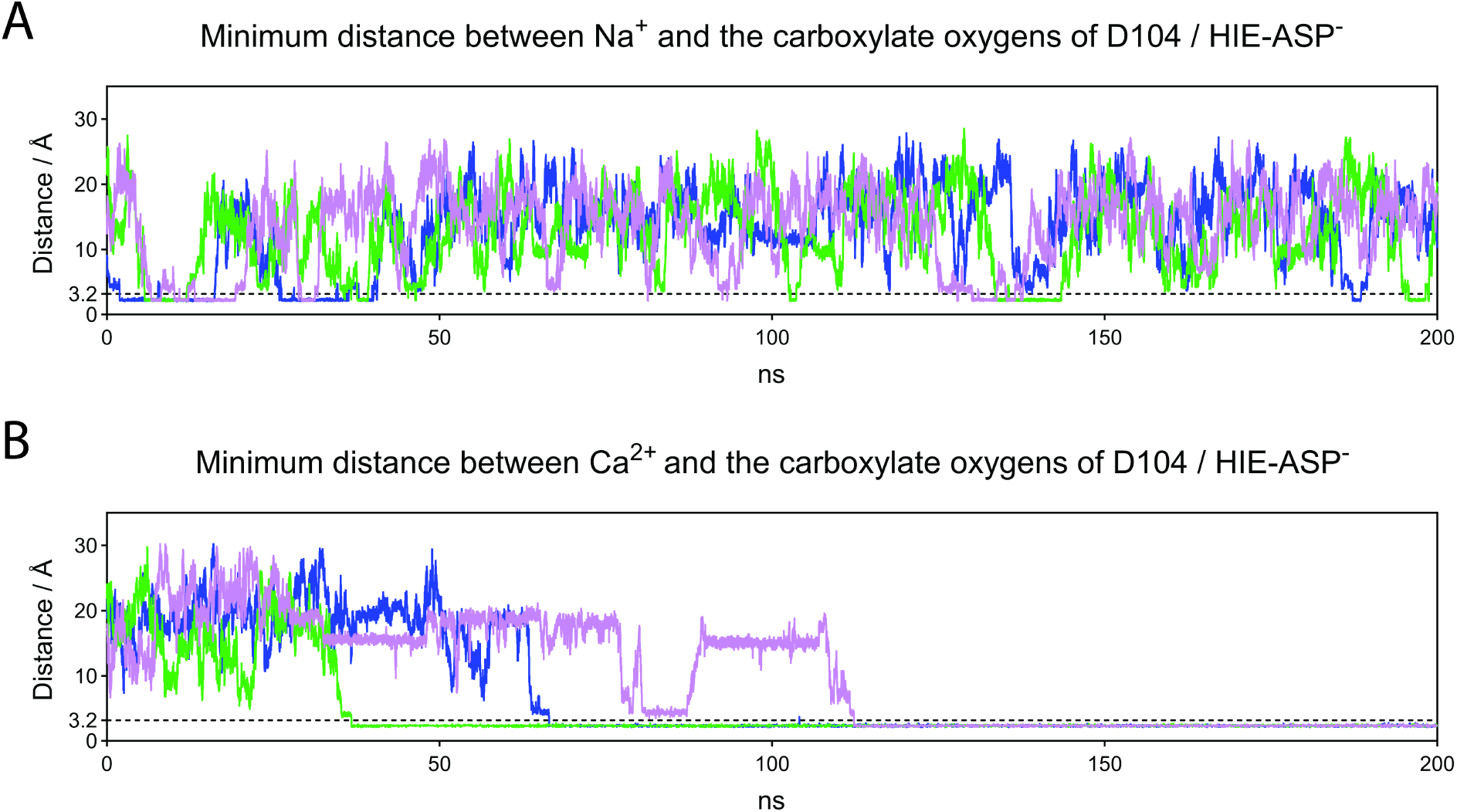
Minimum cation distances to D104 carboxylate oxygens in fixed-protonation conventional MD simulations. Minimum distance between Na⁺ (**A**) or Ca²⁺ (**B**) and either D104 carboxylate oxygen in explicit solvent containing 100 mM NaCl (A) or 50 mM CaCl₂ (B). H102 and D104 were fixed as HIE⁰ and ASP^-^, respectively. Colored traces represent three independent 200-ns simulations. Dashed horizontal lines mark the 3.2 Å coordination cutoff.

**Table S1.** pH-dependent protonation of titratable residues in hen egg-white lysozyme (HEWL). Values at each pH represent the fraction of each residue in its protonated state, as determined by pH-REMD simulations. Apparent p*K*ₐ values and Hill coefficients (*n*_H_) were obtained by fitting the titration curves to the generalized Henderson-Hasselbalch equation. The p*K*ₐ values of D48, D66, D87 and D119 were estimated by extrapolating the fitted curves below the simulated pH range (2.8-7.0). MAE and RMSE denote the mean absolute error and root-mean-square error, respectively, between calculated and experimental p*K*ₐ values. Experimental p*K*ₐ values (*) were taken from Webb et al. (2011).

| **pH** | **E7** | **H15** | **D18** | **E35** | **D48** | **D52** | **D66** | **D87** | **D101** | **D119** |
|---|---|---|---|---|---|---|---|---|---|---|
| 2.8 | 0.79692 | 0.9981 | 0.60546 | 0.99987 | 0.39995 | 0.75497 | 0.31766 | 0.23044 | 0.98824 | 0.41291 |
| 3.4 | 0.56307 | 0.99609 | 0.3322 | 0.999 | 0.22198 | 0.47779 | 0.13002 | 0.08604 | 0.96871 | 0.19962 |
| 4 | 0.29006 | 0.99069 | 0.12583 | 0.9952 | 0.08047 | 0.23281 | 0.04291 | 0.0272 | 0.90176 | 0.07557 |
| 4.6 | 0.1108 | 0.97294 | 0.0391 | 0.98407 | 0.02521 | 0.08319 | 0.01185 | 0.01094 | 0.71602 | 0.02691 |
| 5.2 | 0.03751 | 0.9208 | 0.01046 | 0.9377 | 0.00739 | 0.02364 | 0.00428 | 0.00436 | 0.39878 | 0.0077 |
| 5.8 | 0.01262 | 0.77702 | 0.00299 | 0.79664 | 0.00181 | 0.00699 | 0.00098 | 0.0027 | 0.15368 | 0.00152 |
| 6.4 | 0.00367 | 0.47183 | 0.0007 | 0.50339 | 0.00056 | 0.00256 | 0 | 0.00192 | 0.0586 | 0.00056 |
| 7 | 0.00141 | 0.1693 | 0.00026 | 0.19074 | 7.2E-05 | 0.00102 | 0 | 0.0006 | 0.02319 | 0.00026 |
| pka | 3.53 | 6.34 | 3.03 | 6.4 | 2.57 | 3.37 | 2.41 | 2.18 | 5.02 | 2.61 |
| nH | 0.83 | 1 | 0.85 | 1.01 | 0.71 | 0.85 | 0.84 | 0.84 | 0.94 | 0.77 |
| Experimental pKa* | 2.6 | 5.5 | 2.8 | 6.1 | 1.4 | 3.6 | 1.2 | 2.2 | 4.5 | 3.5 |
| RAE | 0.6332 |  |  |  |  |  |  |  |  |  |
| RMSE | 0.7512 |  |  |  |  |  |  |  |  |  |

**Table S2.**
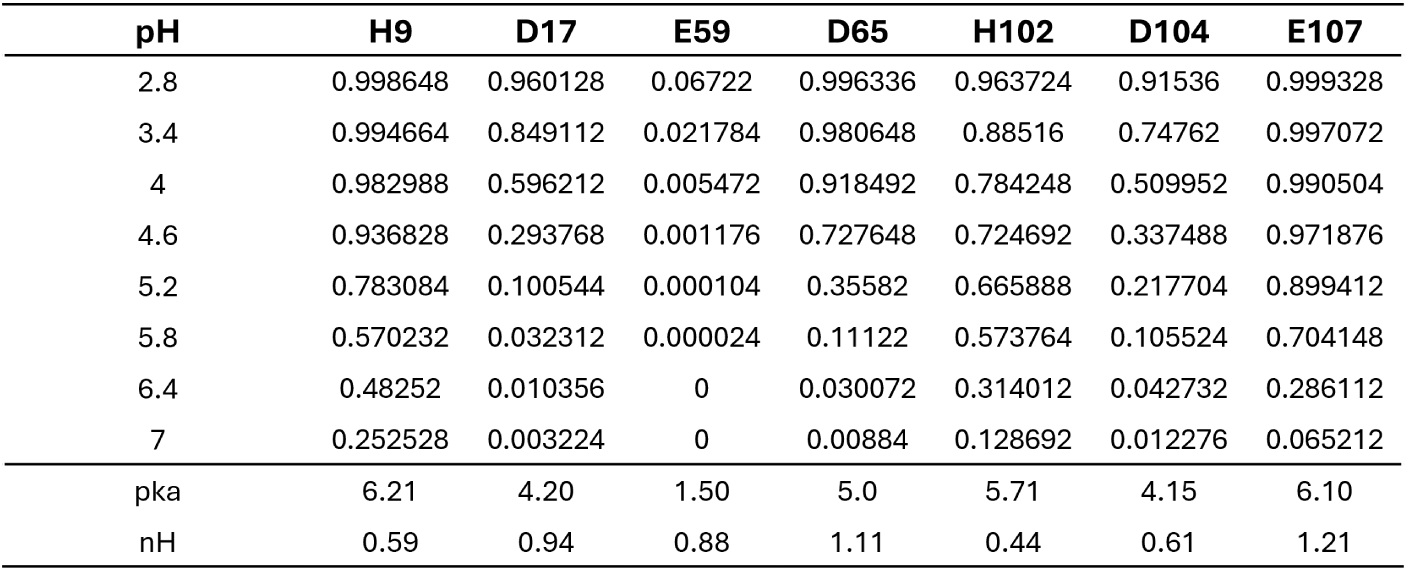
pH-dependent protonation of titratable residues in domain-1 of CsEXPA1. Values at each pH represent the fraction of each residue in its protonated state, as determined by pH-REMD simulations. Apparent p*K*ₐ values and Hill coefficients (*n*_H_) were obtained by fitting the titration curves to the generalized Henderson-Hasselbalch equation. The p*K*ₐ of E59 (1.50) was estimated by extrapolating the fitted curve below the simulated pH range (2.8-7.0).

| **pH** | **H9** | **D17** | **E59** | **D65** | **H102** | **D104** | **E107** |
|---|---|---|---|---|---|---|---|
| 2.8 | 0.998648 | 0.960128 | 0.06722 | 0.996336 | 0.963724 | 0.91536 | 0.999328 |
| 3.4 | 0.994664 | 0.849112 | 0.021784 | 0.980648 | 0.88516 | 0.74762 | 0.997072 |
| 4 | 0.982988 | 0.596212 | 0.005472 | 0.918492 | 0.784248 | 0.509952 | 0.990504 |
| 4.6 | 0.936828 | 0.293768 | 0.001176 | 0.727648 | 0.724692 | 0.337488 | 0.971876 |
| 5.2 | 0.783084 | 0.100544 | 0.000104 | 0.35582 | 0.665888 | 0.217704 | 0.899412 |
| 5.8 | 0.570232 | 0.032312 | 0.000024 | 0.11122 | 0.573764 | 0.105524 | 0.704148 |
| 6.4 | 0.48252 | 0.010356 | 0 | 0.030072 | 0.314012 | 0.042732 | 0.286112 |
| 7 | 0.252528 | 0.003224 | 0 | 0.00884 | 0.128692 | 0.012276 | 0.065212 |
| pka | 6.21 | 4.20 | 1.50 | 5.0 | 5.71 | 4.15 | 6.10 |
| nH | 0.59 | 0.94 | 0.88 | 1.11 | 0.44 | 0.61 | 1.21 |

**Table S3.** pH-dependent protonation of titratable residues in domain-1 of CsEXPA1 with 100 mM NaCl in the explicit solvent. Values at each pH represent the fraction of each residue in its protonated state, as determined by pH-REMD simulations. Apparent p*K*ₐ values and Hill coefficients (*n*_H_) were obtained by fitting the titration curves to the generalized Henderson-Hasselbalch equation. The p*K*ₐ of E59 (1.10) was estimated by extrapolating the fitted curve below the simulated pH range (2.8-7.0).

| **pH** | **H9** | **D17** | **E59** | **D65** | **H102** | **D104** | **E107** |
|---|---|---|---|---|---|---|---|
| 2.8 | 0.99784 | 0.947792 | 0.033244 | 0.989332 | 0.981492 | 0.822024 | 0.996664 |
| 3.4 | 0.993376 | 0.823612 | 0.010484 | 0.961052 | 0.946188 | 0.546192 | 0.993164 |
| 4 | 0.980492 | 0.564284 | 0.003148 | 0.855684 | 0.904112 | 0.268424 | 0.988792 |
| 4.6 | 0.944688 | 0.2707 | 0.000684 | 0.626748 | 0.826144 | 0.120968 | 0.980048 |
| 5.2 | 0.866136 | 0.097608 | 0.000288 | 0.323276 | 0.625888 | 0.061288 | 0.943404 |
| 5.8 | 0.767988 | 0.03528 | 0.000084 | 0.103564 | 0.359544 | 0.031692 | 0.789336 |
| 6.4 | 0.61784 | 0.011832 | 0.000036 | 0.023472 | 0.189752 | 0.025764 | 0.38214 |
| 7 | 0.288608 | 0.004652 | 0 | 0.007372 | 0.0777 | 0.008412 | 0.091688 |
| pka | 6.56 | 4.13 | 1.10 | 4.84 | 5.49 | 3.52 | 6.24 |
| nH | 0.7 | 0.91 | 0.86 | 0.94 | 0.71 | 0.84 | 1.28 |

## Materials and Methods

### Collection of cucumber hypocotyls and maize coleoptiles and measurement of extension

Cucumber (Cucumis sativus) seeds were sown on germination paper moistened with distilled water and grown in darkness at 26 °C for 80 h, until seedlings reached 2.5-3 cm in length. Apical hypocotyl segments were harvested and stored at -80 °C until use. After thawing, the cuticle was gently abraded to facilitate buffer penetration. The freeze-thawed hypocotyls were placed between two glass slides and compressed under a 750-g load for 20 min to remove residual liquid. A 0.5- cm apical segment was then clamped at both ends and subjected to a 20-g constant load. Segment length was recorded every 30 s, and extension rate were calculated as described previously (*4*). To inactivate endogenous α-expansin activity, compressed hypocotyls sandwiched between glass slides were immersed in boiling distilled water for 60 s and then immediately cooled in fresh distilled water.

Maize (Zea mays) seeds were sown on germination paper moistened with distilled water and grown in darkness at 26 °C for 80 h, until coleoptiles reached 2.5-3 cm in length. Coleoptiles were harvested, and the cuticle was gently abraded to facilitate buffer penetration. They were then stored at -80 °C until used in extension assays.

### Tensile testing and stiffness analysis of hypocotyl walls

Freeze-thawed cucumber hypocotyl wall specimens were incubated for 45 min in 20 mM HEPES buffer (pH 7.0) containing 0 or 100 mM NaCl. Specimens were stretched along the growth axis using an Instron 68SC-05 universal testing machine, with an initial gauge length of 5 mm and a constant crosshead speed of 3 mm/min, to 50% strain or until specimen failure, whichever occurred first. Force and extension were recorded simultaneously. Extension was zeroed at the first recorded point at which the force reached or exceeded 0.1 gf. Percentage strain was calculated as,

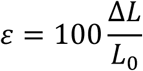

where *ΔL* is extension from this reference point and *L_0_* = 5 mm.

Local apparent axial tensile stiffness was estimated at 5% increments from 5% to 40% strain. For each specimen, a fitting window comprised 101 consecutive force-strain data pairs: the recorded point closest to the target strain and the 50 immediately preceding and 50 immediately following points. Each window spanned approximately ±1 percentage point of strain. Force was fitted against percentage strain by ordinary least-squares linear regression with an intercept,

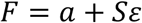

where *a* is the fitted intercept. The local slope, *S*, was calculated as

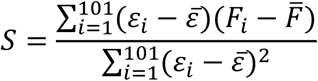

where *ɛ̅* and *F̅* are the mean percentage strain and force within the fitting window, respectively. Slopes were calculated separately for each specimen and expressed in gf per percentage point of strain.

### Crude *α*-expansin extraction

Crude α-expansin extraction from cucumber hypocotyl has been detailed previously, and the cell wall loosening activity from the extraction has been demonstrated to originate from α-expansin (*4*). Briefly, fresh cucumber hypocotyls germinating in the dark at 26°C for 80 hours were homogenized, and the cell wall fragments were collected and washed. Then, crude α-expansin was solubilized in 1 M NaCl and then precipitated with saturated (NH_4_)_2_SO_4_. To assess the protein content, the crude protein diluted in acidic buffer for the acid growth assay was centrifuged at 12000 rpm for 5 minutes at 4 °C, and the total protein concentration was immediately determined to 360 μg/ml by measuring the absorbance at 280 nm.

### Arabidopsis hypocotyl elongation assay

Arabidopsis seedlings were grown in the dark at 22°C for 3 days. Etiolated hypocotyls were immobilized on 0.5% agarose pads containing 20 mM MES buffer, pH 5.2, and imaged under a microscope every 2 min. Hypocotyl length was quantified over 30 min before treatment, during treatment, and after washout. For treatment, seedlings were transferred to 0.5% agarose pads containing 20 mM MES, pH 5.2, supplemented with either 20 mM NaCl or 40 mM mannitol. For recovery, seedlings were rinsed and transferred back to agarose pads containing 20 mM MES, pH 5.2, without added solute.3

### Molecular dynamics simulations

The CsEXPA1 protein structure was obtained from the AlphaFold Protein Structure Database (AF-Q39625-F1-v6)(*37, 38*). Molecular topologies were generated using tleap in AmberTools(*58*). For pH-REMD simulations, the protein was parameterized with the Amber ff19SB force field, and water with the OPC model; constant-pH-compatible parameters were loaded using leaprc.constph. For fixed-protonation conventional MD simulations, the same ff19SB, and OPC parameter sets were used. For all systems, OPC-matched Li-Merz ion parameters were loaded using frcmod.ionslm_126_opc (Na^+^) and frcmod.ionslm_1264_opc (Ca^2+^). Prepared PDB structures were loaded into tleap, and four predicted disulfide bonds were explicitly assigned between Cys26-Cys54, Cys57-Cys130, Cys62-Cys69, and Cys82-Cys96. Systems were checked, solvated in a truncated octahedral OPC water box with a minimum 12 Å buffer, and neutralized with Cl^-^ counterions using addIons2. Explicit Na⁺/Cl^-^ or Ca²⁺/Cl^-^ ions were added to produce final NaCl and CaCl₂ concentrations of 100 mM and 50 mM, respectively.

All molecular dynamics simulations were performed with Amber22 (*59*).

pH-REMD simulations followed the hybrid-solvent discrete-protonation protocol of Swails et al. (2014)(*47*), with modifications described below. Protein conformations were sampled by MD in explicit solvent, whereas discrete protonation-state changes were evaluated using Monte Carlo (MC) moves with a generalized Born (GB) implicit-solvent model. Replicas spanned pH 2.8-7.0 in 0.6-unit increments. Each replica was first equilibrated for 2 ns under NVT conditions at its assigned pH with constant-pH sampling enabled. Production simulations were then performed for 50 ns per replica, with exchange attempts between neighboring pH replicas every 1,000 MD steps. Protonation-state updates were attempted every 100 MD steps using the hybrid-solvent constant-pH implementation (icnstph = 2). During each update, explicit water and nonstructural ions were temporarily excluded from the energy evaluation, and proposed protonation states were assessed by MC using the GB model at fixed solute coordinates and the assigned pH. The implicit salt concentration used in the GB calculations was set to 0.1M. Explicit solvent and ions were subsequently restored. Following accepted protonation-state changes, the solvent and ions were relaxed for 100 MD steps with the solute held fixed before full explicit-solvent MD resumed. Simulations were carried out at 300 K using Langevin dynamics with a collision frequency of 2.0 ps⁻¹ and automatically assigned random seeds (ig = -1). Bonds involving hydrogen atoms were constrained with SHAKE, allowing a 2-fs time step. Periodic boundary conditions were applied with a 10 Å nonbonded cutoff. Coordinates were saved every 10 ps and restart files every 100 ps. Convergence was assessed by comparing protonation fractions across successive 10-ns blocks. For protocol validation, simulations of hen egg-white lysozyme (HEWL) were initiated from its crystal structure obtained from the Protein Data Bank (PDB ID: 1AKI).

For each titratable residue, the protonated fraction at each pH was calculated as:

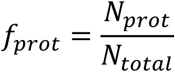

where *N_prot_* and *N_total_* denote the number of protonated samples and the total number of samples collected at that pH, respectively. Protonated microstates were combined for Asp and Glu. For His, the protonated fraction corresponded to HIP^+^, whereas HID⁰ and HIE⁰ together constituted the neutral fraction. Apparent p*K*ₐ values and Hill coefficients (*n*_H_) were obtained by nonlinear least-squares fitting to the generalized Henderson-Hasselbalch (Hill) equation:

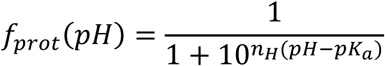

The fitted p*K*ₐ corresponds to the pH at which *f*_prot_ = 0.5, and *n*_H_ describes the steepness of the titration transition. A value of *n*_H_ = 1 gives standard Henderson-Hasselbalch behavior, whereas values below or above 1 indicate broader or steeper transitions, respectively.

For conventional MD simulations, the protonation states of H102 and D104 were assigned as indicated. All other titratable residues were kept in their default protonation states: Asp as ASP^-^, Glu as GLU^-^, and His as HIE^0^. All quantitative analyses were based on three independent replicas. Each replica was first equilibrated for 1 ns under NPT conditions and then subjected to 500 ns of production MD. Replicas used time-dependent Langevin random seeds (ig=-1), and their initial velocities were verified to be distinct. Production simulations were carried out under NPT conditions using the Monte Carlo Barostat. Simulations were performed at 300 K with Langevin dynamics and a collision frequency of 2.0 ps^-^¹. Bonds involving hydrogen atoms were constrained with SHAKE, allowing a 2-fs time step. Nonbonded interactions were calculated with a 10 Å cutoff under periodic boundary conditions. Coordinates were saved every 10 ps, and restart files were written every 100 ps. All trajectory analyses were performed using custom Python scripts with NumPy and Amber NetCDF/topology files, applying periodic minimum-image corrections for distance-based measurements.

Hydrogen bonds were identified using geometric criteria. A hydrogen bond was assigned when the donor-acceptor distance was ≤ 3.2 Å and the donor-hydrogen-acceptor angle was ≥ 120°. Salt bridges were defined by a minimum distance ≤ 4.0 Å between oppositely charged heavy atoms, including H102 imidazolium nitrogens and D104 carboxylate oxygens in the HIP⁺-ASP^-^ state. Occupancies were calculated as the fraction of trajectory frames satisfying the corresponding criterion, with periodic minimum-image corrections applied to all distance measurements.

